# Uncovering the signaling networks of disseminated glioblastoma cells *in vivo* with INSIGHT

**DOI:** 10.1101/2024.12.04.626717

**Authors:** Ryuhjin Ahn, Alicia D. D’Souza, Lawrence Long, Yufei Cui, Danielle Burgenske, Katrina K. Bakken, Lauren L. Ott, Brett L. Carlson, Grace Zhou, Tomer M. Yaron-Barir, Ishwar N. Kohale, Charles A. Whittaker, Cameron T. Flower, Jeffrey Wyckoff, Ann Tuma, Jared L. Johnson, Jann Sarkaria, Forest M. White

## Abstract

Dysregulation of intracellular signaling networks underpins cancer. However, a systems-level elucidation of how signaling networks within distinct cell subpopulations drive cancer progression *in vivo* has been unattainable due to technical limitations. We developed INSIGHT (INvestigating SIGnaling network of specific cell subpopulation in Heterogeneous Tissue), a new platform technology combining fluorescence-activated cell sorting with ultra-sensitive mass spectrometry to enable phosphoproteomic characterization of rare and discrete cell subpopulations from fixed tissues. We demonstrated the broad utility of INSIGHT by analyzing the oligodendroglial cell-specific signaling network in the mouse brain. We then applied INSIGHT to investigate the rare, disseminated tumor cell subpopulation in glioblastoma patient-derived xenograft models. INSIGHT uncovered a global rewiring of signaling networks with tumor cell dissemination, marked by a transition from proliferation-associated signaling in the primary tumor cells to signaling associated with postsynapse, neuronal migration, and ion homeostasis in disseminated tumor cells. We reveal interconnections between signaling circuitries within the networks, with numerous proteins, including GluA2, exhibiting altered phosphorylation without protein expression changes, emphasizing the role of post-translational modifications in glioblastoma dissemination. We validated key phosphorylation changes and inferred differentially active kinases with tumor spread to offer new systems-level insights into glioblastoma dissemination mechanisms *in vivo*. INSIGHT is generally applicable to a wide range of biological systems without genetic engineering and provides quantitative phosphorylation and protein expression data for selected cell subpopulations from heterogeneous tissues.

## Introduction

Dysregulation of intracellular signaling networks drives cancer^1,2^. Cellular signaling is mediated by dynamic phosphorylation and dephosphorylation of proteins by kinases and phosphatases primarily on tyrosine, serine, and threonine residues, which directly modulate protein function^3–5^. The significance of phosphorylation in cancer is underscored by kinase inhibitors representing one of the largest classes of FDA-approved cancer drugs^6^. Currently, the systems-level analysis of phosphorylated proteins, or phosphoproteomics, is best achieved by mass spectrometry. However, due to the rapid dynamics of phosphorylation^7^, phosphoproteomics requires rapid sample processing^8,9^, which restricts the analysis to bulk tissues. This obscures the signaling networks originating from rare and diverse cell subpopulations, leaving a substantial blind spot in our understanding of heterogeneous cellular functions and intercellular communications that collectively shape the biological response *in vivo*. In particular, the low cellular stoichiometry of tyrosine phosphorylation (<1% of the phosphoproteome^10^), requires a large sample input (1-15mg^11,12^) for informative analysis, preventing in-depth phosphotyrosine phosphoproteomic analysis at the resolution of single-cell or low numbers of cells *in vivo*. Phosphoprotein-specific antibody-based approaches like multiplexed tissue imaging^13^ do not allow unbiased discovery of signaling networks. Thus, these technical barriers have impeded the systems-level elucidation of how signaling networks of distinct cell subpopulations contribute to cancer progression *in vivo*.

Glioblastoma (GBM) remains a lethal cancer, with recurrence driven by invasive subpopulations of tumor cells that disseminate brain tissue and evade surgical resection, rendering even drastic interventions like hemispherectomy ineffective^14^. GBM cells migrate along anatomical structures including white matter tracts as singlets or clusters^15^. Remarkably, preclinical GBM patient-derived xenograft (PDX) tumor cells retain this behavior, with rare, discrete tumor subpopulations disseminating throughout the mouse brain^16^. Seminal studies have established that GBM cells exploit both neuronal program-dependent and independent mechanisms to disseminate^17–22^. Yet, we lack a systems-level understanding of how the signaling network of GBM orchestrates the dissemination *in vivo*, how its signaling circuitries interconnect, and which of its proteins are regulated by phosphorylation with which kinases, hindering us from comprehensively exposing the therapeutic vulnerabilities of GBM.

To enable the quantification of signaling networks in specific cell subpopulations within complex tissues, including that of rare, disseminated tumor cells that underlie GBM recurrence, we developed INSIGHT (INvestigating SIGnaling network of specific cell subpopulation in Heterogeneous Tissue), a new discovery-based phosphoproteomic and protein expression analysis platform technology that combines ultra-sensitive liquid chromatography-tandem mass spectrometry (LC-MS/MS) and targeted fluorescence-activated cell sorting (FACS) with broad applicability in cancer and beyond. In INSIGHT, physiological phosphorylation sites are preserved by freezing and subsequently fixing before dissociation and cell sorting, enabling systems-level characterization of signaling networks within rare or discrete targeted cell subpopulations. We first demonstrated the utility of INSIGHT by analyzing the oligodendroglial cell-specific signaling network in the mouse brain. We then applied INSIGHT to invasive GBM PDX models, and quantified thousands of phosphoproteins and proteins that compose the signaling network of rare, disseminated GBM subpopulations. We captured interconnections between their diverse signaling circuitries, validated key phosphoproteins altered during dissemination using microscopy, and inferred their active kinases to reveal possible molecular mechanisms underpinning GBM dissemination *in vivo*.

## Results

### Phosphoproteome is preserved by fixation and remains stable through enzymatic dissociation to enable antibody- or fluorescence tag-based cell sorting

To enable the signaling network analysis of discrete cell populations of interest *in vivo*, we established INSIGHT, a platform technology that preserves the unstable phosphoproteome throughout the analysis^7–9^. INSIGHT uses frozen tissue, which is cryosectioned at -20°C, fixed with a commercial fixative, and enzymatically and physically dissociated into single cells to allow the staining and sorting of specific cell types before phosphoproteomic and protein expression analyses by ultra-sensitive LC-MS/MS (**Fig. 1**). To establish this workflow, we first evaluated the effectiveness of fixation in preserving physiological phosphorylation sites compared to the rapid standard sample processing workflow for LC-MS/MS and conventional tissue dissociation conditions. Frozen brain tissue sections were subjected to five distinct conditions: (1) 37°C for 2 hours, (2) 37°C for 1 hour, (3) cold PBS rinse, (4) 10% buffered formalin fixation, and (5) Fixative 1 (Cytofast; Biolegend) fixation (**Extended Data Fig. 1a**). The 37°C incubation, common in single-cell tissue studies^23^, significantly altered protein expression and phosphorylation compared to the control, while both fixation methods more effectively preserved the phosphoproteome. Next, to find the most effective fixative, we benchmarked 10% buffered formalin and Fixative 1 to four other fixatives (**Extended Data Fig. 1b**). We observed that Fixative 1 showed superior identification of antibody-targeted cells and recovery during dissociation while effectively preserving the phosphoproteome (**Extended Data Fig. 1a-b**). Therefore, Fixative 1 was selected as the fixative for the rest of this study.

**Figure 1.**
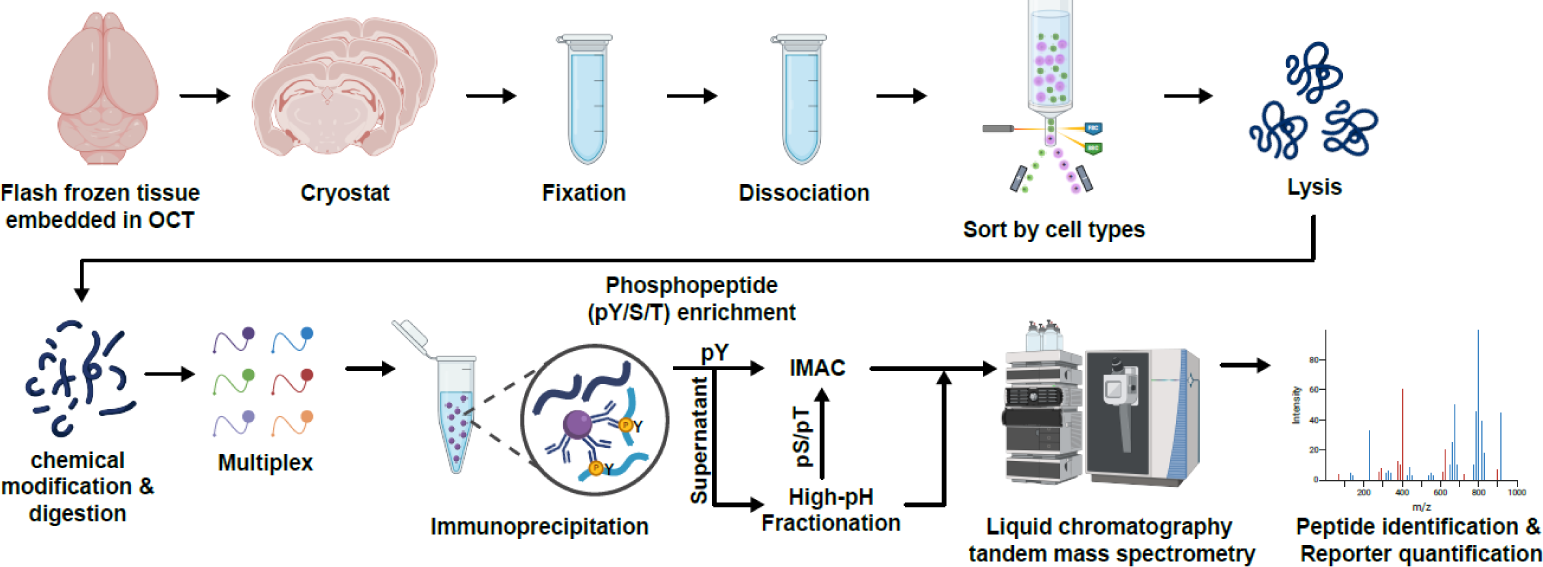
INSIGHT enables cell subpopulation-specific signaling network analysis *in vivo*. Schematic of INSIGHT platform technology. Flash-frozen tissue is cryosectioned, fixed, and dissociated to obtain single cells. Cell subpopulations are sorted by fluorescence-activated cell sorting (FACS) using fluorophore-conjugated antibodies or endogenously expressed fluorophores. Sorted cells are lysed, proteins are digested, and peptides are labeled using tandem mass tags for multiplexed analysis. Phosphotyrosine-containing peptides (pY) are enriched through immunoprecipitation and immobilized metal affinity chromatography (IMAC). Phosphoserine (pS), and phosphothreonine (pT) from pY-IP supernatant undergo high-pH fractionation and phosphopeptide enrichment by IMAC. Both the enriched phosphopeptides and fractionated supernatants are analyzed using liquid chromatography-tandem mass spectrometry (LC-MS/MS) to quantify phosphoproteins and proteins.

Next, to dissociate the fixed tissues for downstream cell-type sorting, we evaluated the impact of enzymatic dissociation on the phosphoproteome and whole proteome of fixed tissues to ensure that they do not degrade peptides before LC-MS/MS analysis. After testing several collagenases, correlation analysis showed that collagenase III best preserved the fixed phosphopeptides (**Extended Data Fig. 2a**), leading us to use it for the rest of this study. To assess the integrity of the cells and the protein loss from INSIGHT-induced membrane permeabilization or disruption, the molecular weight distribution of proteins and phosphoproteins quantified by MS in dissociated and undissociated cells was analyzed (**Extended Data Fig. 2b**). No significant differences were found, indicating that any depletion was likely not due to the loss of smaller proteins from INSIGHT-induced membrane permeabilization or leakiness. We also quantified cell yields from the mouse liver, spleen, and brain, demonstrating the potential applicability of INSIGHT across different tissues (**Extended Data Fig. 2c**).

To confirm that epitopes remain intact for cell isolation by antibody-based sorting and that INSIGHT does not cause nonspecific binding or background signals during flow cytometric analysis, we tested the approach on intracranially injected, EGFR-amplified, TdTomato-tagged GBM PDX tumors (**Extended Data Fig. 2d**). Anti-human EGFR and anti-HLA-A/B/C antibodies specifically bound the TdTomato+ cells in the dissociated PDX-bearing mouse brain, confirming the preservation of surface protein epitopes with no detectable unspecific binding (**Extended Data Fig. 2e**). This result also highlights that fluorescently tagged cells may be subjected to INSIGHT to characterize their phosphoproteome and protein expression, broadening the applicability of INSIGHT in genetically engineered model organisms where specific populations can be fluorescently tagged.

### INSIGHT can be used to map cell type-specific signaling networks *in vivo*

As a proof of principle to demonstrate that INSIGHT can characterize the signaling networks of specific cell subpopulations from tissues, we focused on oligodendrocyte lineage cells of the brain. Oligodendrocyte lineage cells (oligodendrocytes and oligodendrocyte progenitor cells), serve critical roles in myelination and provide metabolic support to neurons and tumors^24,25^. We leveraged the cell-specific marker O4 antigen to isolate O4-positive oligodendrocyte lineage cells and O4-negative cells for analysis by INSIGHT (**Fig. 2a, b**). This resulted in the identification of 4,428 unique proteins and 12,465 phosphopeptides at a 1% FDR threshold. To ensure high-quality phosphopeptide and protein identification, peptide spectral matches (PSMs) were filtered using a high-stringency (HS) ion score cutoff of 20, established through extensive manual validation of spectra^26^. This refinement yielded 4,269 proteins and 9,351 phosphopeptides (**Fig. 2c**), comprising 9,767 phosphoserine (pS), 1,780 phosphothreonine (pT), and 673 phosphotyrosine (pY) (**Fig. 2d**). Analysis of the O4-positive proteome against the Tabula Muris dataset showed oligodendrocyte enrichment (**Fig. 2e**), and oligodendrocyte marker proteins were specifically enriched in O4-positive proteome and phosphoproteome (**Fig. 2f-g**). Moreover, O4-positive cells were enriched in oligodendroglial processes, while O4-negative cells were enriched in neuronal processes (**Fig. 2h**). Collectively, these results confirmed that INSIGHT successfully profiled the signaling network of enriched oligodendrocyte lineage cells from fixed brain tissues. Furthermore, INSIGHT mapped O4-positive and negative cell-specific expression of kinases, phosphatases, and their phosphorylated forms, revealing their expression patterns and potential activation status (**Fig. 2i**). Next, leveraging cell-type specific protein expression data, we used NicheNet^27^ to infer ligand-receptor interactions between O4-negative and O4-positive cells that may regulate myelin maintenance within O4-positive cells. We then integrated ligand-receptor interaction predictions with receptor phosphorylation data to further refine the predicted interaction landscape. This strategy enabled us to infer potential ligand-receptor signaling dynamics across the two subpopulations that contribute to myelin maintenance (**Fig. 2j**), highlighting the potential of INSIGHT in uncovering physiologically relevant cell-cell communication mechanisms at the protein and phosphoprotein-level. Taken together, we validated the specificity of the isolation strategy of INSIGHT by mapping the signaling network of oligodendroglial cells and demonstrated the utility of INSIGHT in inferring physiologically relevant information *in vivo*.

**Figure 2.**
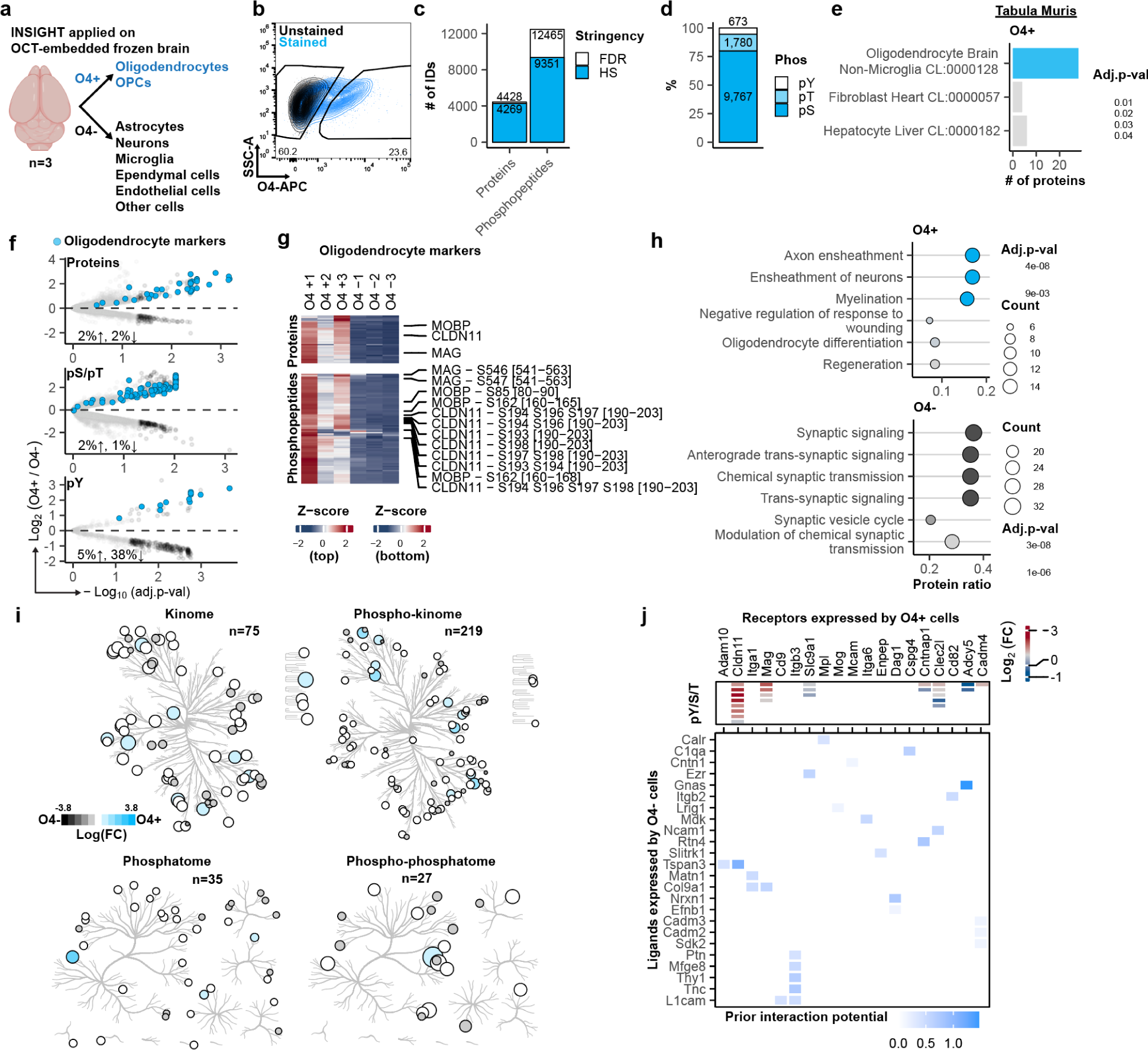
Signaling network of oligodendroglial cells *in vivo*. (a) O4-positive oligodendroglial cells (oligodendrocytes, oligodendrocyte progenitor cells (OPCs)) were sorted from O4-negative cells in three mouse brains. (b) Representative FACS profile of O4-positive and O4-negative population, with y-axis representing side scatter area and x-axis representing O4 fluorescence intensity. (c) The number of identified proteins and phosphopeptides (combined output from phosphotyrosine immunoprecipitation and global phosphoproteomics) at 1% FDR and in-house determined highly stringent (HS) filter. All the subsequent analyses were done using HS identifications. (d) Bar graph showing the percentages of pY, pS, and pT quantified across phosphopeptides in **c**. (e) Cell-type enrichment analysis using the Tabula Muris database using O4-positive specific proteins (fold change > 1 and adj. p-val ≤ 0.1) relative to O4-negative specific proteins. (f) Volcano plots showing differentially expressed pY, pS, pT and proteins between O4-positive and O4-negative cells. The top quadrant represents IDs enriched in O4-positive cells compared to O4-negative cells. Oligodendrocyte markers were overlaid (blue). Differentially expressed IDs with 1.5-fold change and adj. p-val ≤ 0.05 were colored in black and their percentages are shown at the bottom left. (g) Heatmap depicting all the oligodendrocyte markers quantified at the protein (upper) and phosphopeptide (pY/pS/pT) levels (lower) shown in f. Z-scores indicate relative expression levels (red = higher, blue = lower). Select markers are shown as examples. (h) Enriched biological processes in the proteome of O4-positive and O4-negative cells (1.5-fold differentially expressed proteins at adj. p-val ≤ 0.05). The top annotations are shown. The count represents the number of genes matched. (i) Kinome and phosphatome tree depicting the level of O4-positive and O4-negative enriched total and phosphorylated kinases and phosphatases. Node sizes are based on the log-transformed fold change of O4-positive to O4-negative expression data. (j) Heatmap depicting the prior interaction potential between the receptors expressed by O4-positive cells and the ligands expressed by O4-negative cells for myelin maintenance as assessed by NicheNet. The relative abundance of pY, pS, and pT residues between O4-positive and O4− cells, quantified by MS (Log_2_ fold-change), for receptors identified in O4-positive cells is presented (top).

### Signaling network of rare, disseminated glioblastoma cells is uncovered by INSIGHT

Having demonstrated the feasibility of INSIGHT for cell subpopulation-specific phosphoproteomics, we leveraged INSIGHT to characterize the signaling networks of invasive, disseminated epidermal growth factor receptor (EGFR)-amplified GBM cells *in vivo*. EGFR overexpression, in its wild-type or constitutively active EGFRvIII mutant form, is found in over 40% of GBM patients and drives tumorigenicity^28,29^. We used two PDX models, GBM6 (G6) and GBM12 (G12), derived from untreated primary IDH-wildtype GBM^30^. G12 harbors wild-type EGFR amplification, whereas G6 carries an EGFRvIII mutation^31,32^. Implantation of PDX tumors into the right hemispheres of mice led to contralateral tumor dissemination via white matter tracks of the corpus callosum as seen in patients^33^ (**Fig. 3a**). Notably, G6 primary tumor cells showed greater tumor spread, more diffuse boundaries, and higher normal cell content compared to G12 primary tumor cells at the coronal section with the highest primary tumor burden (**Fig. 3a, Extended Data Fig. 3a, b**), indicating a more invasive phenotype. We then set out to isolate and analyze the tumor cells from each hemisphere by enriching for human EGFR-expressing cells (**Fig. 3b**). To do so, G6 and G12 PDX tumor-bearing brains were resected at moribund, flash-frozen, and cut down at the mid-sagittal plane. Each hemisphere was cryosectioned, fixed, and dissociated to yield single cells (**Fig. 3c**). Cells were then stained with a human-specific anti-EGFR antibody (also detects EGFRvIII) and sorted to isolate EGFR-positive rare, disseminated tumor cells (<1%) from the left hemisphere and EGFR-positive primary tumor cells from the right hemisphere (**Fig. 3b, d, e, Extended Data Fig. 3c, d**). The higher number of disseminated cells in G6 compared to G12 corroborated the co-immunofluorescence (co-IF) staining result (**Fig. 3a, e, Extended Data Fig. 3a, b**). Additionally, EGFR-negative cells (non-malignant cells; “normal cells”) and bulk unsorted mixtures of cells were taken from each hemisphere for comparison. In total, we analyzed nine G6 tumor-bearing brains and three G12 tumor-bearing brains, all of which were subjected to phosphoproteomic and protein expression analysis via LC-MS/MS (**Fig. 3b**).

**Figure 3.**
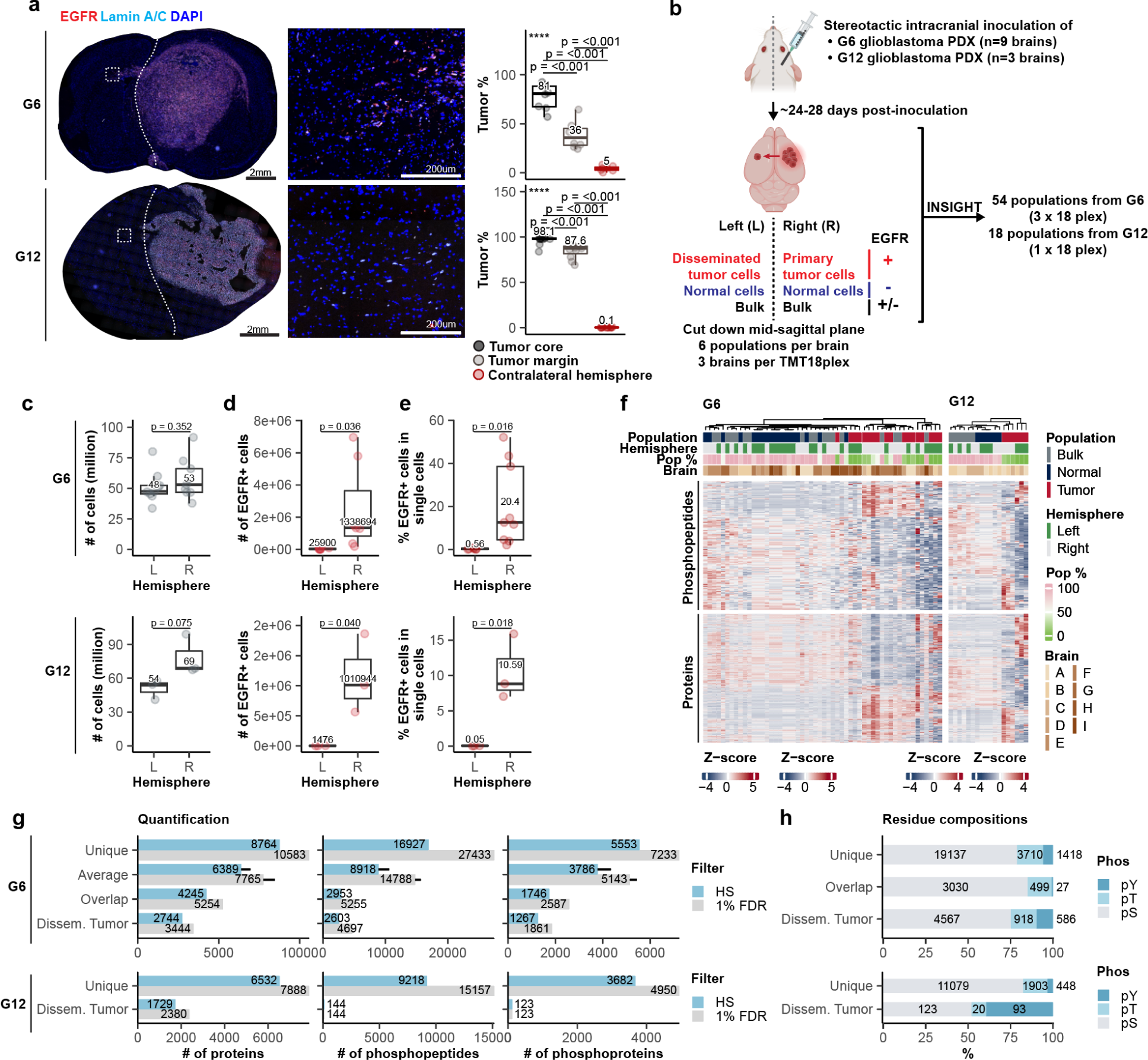
Application of INSIGHT on GBM PDX models. (a) Representative images from G6 and G12 GBM PDX-bearing brains stained for human-specific EGFR, human-specific Lamin A/C, and DAPI (nuclei) in coronal sections with the highest primary tumor burden. White dotted lines mark the midline of the brain, separating the left and right cerebral hemispheres. The middle panel shows the magnified images of the region marked by the dotted rectangle in the left panel. The percentage of tumor cells (EGFR+ Lamin A/C+ /DAPI+) was quantified based on three regions: the core of the primary tumor, the margins of the primary tumor containing the invading tumors, and the entire left hemisphere containing disseminated tumors. Scale bar = 2 mm for the whole sections, 200 µm for the magnified images. (b) Schematic of the experimental setup. EGFRvIII-amplified G6 and wild-type EGFR-amplified G12 GBM PDXs were stereotactically injected into the right hemispheres of mice (n=9 and n=3, respectively). After 24-28 days, some primary tumor cells in the right hemisphere disseminated into the left hemisphere. Brains were removed and flash-frozen, cut down the mid-sagittal plane, and each hemisphere was sectioned, fixed, dissociated, stained with a human-specific anti-EGFR antibody, and sorted by FACS. Samples (EGFR+ tumor cells, EGFR-normal cells, bulk mixture of cells) were processed, multiplexed, and subjected to phosphoproteomic and protein expression analysis by LC-MS/MS. (c) Total cells retrieved from the dissociation of the left and right hemispheres of the G6 and G12 PDX tumor-bearing brains. (d) Number of EGFR+ cells retrieved by FACS. Median values are annotated. (e) Percentage of EGFR+ cells in single cells by FACS. Median values are annotated. (f) Hierarchical clustering of G6 and G12 PDX samples processed by INSIGHT based on identified proteins and phosphopeptides (rows). Only the overlapping, highly stringently (HS) filtered identifications across multiplexed sets are depicted, and any PSMs with missing values were removed except for those identified only in the disseminated tumor (where missing values were imputed to enable comparison; refer to Method). Pop % = percentage of the population relative to the respective total single cells sorted. The order of the heatmap legends (z-score) is G6 phosphopeptide, G6 protein, G12 phosphopeptide, and G12 protein. (g) Number of proteins, phosphopeptides, and phosphoproteins passing the HS and 1% FDR filter across all experiments for G6 and G12 PDX tumor-bearing brains, including unique IDs (unique), shared IDs across the three runs in G6 (overlap; standard deviation shown), and those detected in disseminated tumor cells (dissem. tumor). (h) The percentage of pY, pS, and pT quantified for phosphopeptides that passed the HS filter in **g**. The numbers represent residues from uniquely mapped IDs (unique), shared IDs (overlap), and those detected in disseminated tumor (Dissem. Tumor), across the three runs.

Upon unsupervised hierarchical clustering of the MS data, primary and disseminated tumors clustered together and away from bulk mixtures and normal cells, indicating that the signaling profiles of tumor cells, regardless of location, were similar to each other and distinct from normal cells (**Fig. 3f**). The bulk cells clustered with EGFR-negative normal cells, suggesting that a significant proportion of unsorted cells consisted of normal cells, as expected. Across all G6 samples, we identified 8,764 unique proteins, with an average of 6,389 proteins across plexes and 4,245 proteins detected in all plexes (**Fig. 3g**). In G12, we identified 6,532 unique proteins and 9,218 phosphopeptides (**Fig. 3g**). The HS-filtered data comprised of 24,265 unique phosphosites across 16,927 phosphopeptides mapping to 5,553 proteins across all G6 runs, while in G12, 13,430 unique phosphosites across 9218 phosphopeptides were mapped to 3,682 proteins (**Fig. 3g, h**). When analyzing only those IDs that were expressed in disseminated tumor cells, 2,744 and 1,729 unique proteins were detected in G6 and G12, respectively, and 2,603 and 144 unique phosphopeptides were identified in G6 and G12, respectively (**Fig. 3g**). Thus, despite limited cell numbers, INSIGHT enabled systems-level quantification of phosphopeptides and proteins from both tumor and normal cells in the left and right hemispheres.

### INSIGHT quantifies and differentiates the signaling networks of tumor and normal cells in PDX models to enable the inference of tumor-specific kinase activity

Next, we performed differential expression analysis of the phosphoproteome and proteome between primary tumor cells and normal cells to evaluate whether the INSIGHT workflow could effectively identify the tumor cell-specific signaling network. In both G6 and G12 models pronounced differences were observed between the proteome of tumor cells and normal cells, as expected (**Fig. 4a**, **Extended Data Fig. 4a**). Importantly, murine oligodendrocyte-specific proteins were specifically enriched in the murine normal cell proteome, while human Olig2 was expressed by human primary G6 tumor cells, consistent with a previous report^34^, indicating that INSIGHT successfully separated human tumor cells from the murine normal brain cells, which has been previously impossible in bulk phosphoproteomics of PDX models^35^. Moreover, primary tumor cells of both PDX models upregulated biological processes related to enhanced transcriptional and proliferative activity (**Fig. 4b, Extended Data Fig. 4b**). In contrast, normal cells were enriched in GO terms reflecting metabolic homeostasis and neurobiological functions of brain cells (**Fig. 4b, Extended Data Fig. 4b**).

**Figure 4.**
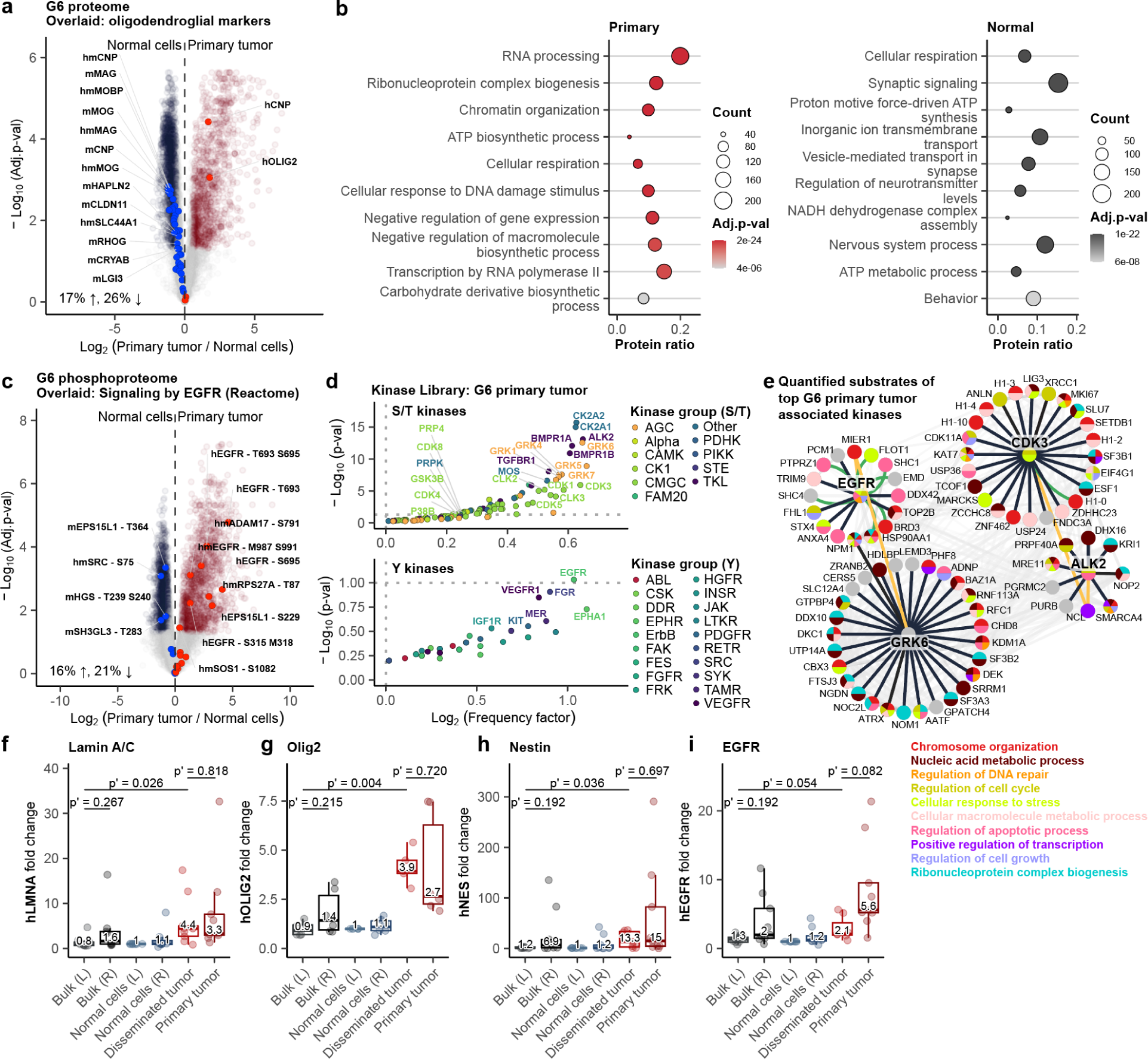
INSIGHT distinguishes the signaling networks of primary tumor cells and normal cells *in vivo*. (a) Volcano plot illustrates differentially enriched proteins between EGFR-positive primary tumor cells and normal cells from the left hemisphere of the G6 PDX model, with oligodendrocyte markers overlaid. The percentages of significantly differentially expressed IDs (1.5-fold change, adj. p-val ≤ 0.05) are shown in the bottom left corner. Lowercase prefixes indicate proteins specific to humans (h), mice (m), or shared between humans and mice (hm). (b) Biological processes significantly enriched (adj. p-val ≤ 0.05) in G6 primary tumor cells (left) and normal cells (right) from the right hemisphere based on their protein expression (1.5-fold change, adj. p-val ≤ 0.05). The top annotations are shown. (c) Volcano plot showing the differentially enriched phosphopeptides in primary tumor cells and normal cells from the right hemisphere of the G6 PDX model. EGFR signaling components based on the Reactome database are labeled. The percentage of significantly differentially expressed IDs (1.5-fold change, adj. p-val ≤ 0.05) is shown in the bottom left corner. Lowercase prefixes indicate proteins specific to human (h), mouse (m), or shared between human and mouse (hm). (d) Phosphopeptides (containing pY, pS, pT) mapped to the human proteome and enriched in the G6 primary tumors compared to normal cells from the right hemisphere (1.5-fold change, adj. p-val ≤ 0.1) were used to infer active kinases based on the Kinase Library^4^. (e) Proteins whose phosphorylations were predicted by Kinase Library to be mediated by top-enriched EGFR, GRK6, ALK2, and CDK3 in **d** were analyzed by STRING. Node colors represent significantly enriched biological processes (adj. p-val ≤0.05) associated with the proteins. Grey edges: known protein-protein associations with edge thickness indicating the strength of evidence. Bold black edges: Kinase Library predicted kinase-substrate relationships. Green edges: predicted kinase-substrate relationships that also had known protein-protein associations by STRING. Yellow edges: proteins that had two separate phosphorylation sites that were regulated by two kinases. (f-i) Protein levels of human (f) Lamin A/C (LMNA), (g) Olig2, (h) Nestin (NES), and (i) EGFR across bulk cell mixtures, normal cells (EGFR-), and EGFR+ tumor cell populations from the left (L) and right (R) hemispheres of the G6 PDX model by INSIGHT. Missing values in cells other than disseminated tumor cells were imputed with the lowest m/z intensity in each PSM at the level of each TMT experiment (refer to Method) to enable comparison. Adjusted p-values of MS data were calculated using the Benjamini-Hochberg correction. Following the concatenation of three G6 TMT experiments, all samples detecting Lamin A/C, Nestin, EGFR (primary n = 9, disseminated n = 7), and Olig2 (primary n = 6, disseminated n = 5) were plotted.

The phosphoproteome of primary tumor cells also significantly differed from that of normal cells in both PDX models (**Fig. 4c**, **Extended Data Fig. 4c**). Still, a large proportion of phosphopeptides were shared between the tumor cells and normal cells in both PDX models based on the percentage of statistically significant up- and down-regulated phosphopeptides, highlighting that their accurate quantification would be impossible without physical separation using INSIGHT. Importantly, phosphorylated EGFR was significantly enriched in tumor cells, consistent with EGFR amplification in both PDX models, underscoring the effectiveness of INSIGHT in isolating tumor-specific phosphoproteome *in vivo* (**Fig. 4c, Extended Data Fig. 4c**). Using the quantified tumor phosphoproteome and the Kinase Library (contains sequence motif specificities of the human kinome^4,5^), the most active kinases in tumor cells based on the prevalence of their phosphorylated substrates were inferred (**Fig. 4d, Extended Data Fig. 4d**). This analysis identified EGFR as top-ranking tyrosine kinase (**Fig. 4d bottom, Extended Data Fig. 4d bottom**), consistent with EGFR amplification in both PDX models. It also revealed additional GBM-associated serine/threonine kinases (e.g., G protein-coupled receptor kinases (GRKs), Cyclin-dependent kinases (CDKs))^36–39^ as well as others that may contribute to GBM progression. Proteins with phosphorylation sites predicted to be mediated by these kinases showed significant enrichment in interconnected, tumor-related biological processes, highlighting extensive overlap in signaling circuitries (**Fig. 4e**). Together, INSIGHT enabled the differentiation and quantification of signaling networks in normal and tumor cells within PDX models which has been previously impossible in phosphoproteomics, thus allowing for the inference of potentially active kinases specific to tumor cells.

### Disseminated tumor cells retain tumor-specific marker expression while downregulating EGFR and pT693-EGFR relative to primary tumor cells

Next, we assessed the expression of previously reported tumor-specific markers^34,40^ to compare the enrichment of disseminated and primary tumor cells. Olig2, Nestin, and Lamin A/C were comparably expressed in both G6 primary and disseminated tumors, confirming marker retention at distant sites and successful tumor cell isolation (**Fig. 4f-h**). Intriguingly, EGFR expression (mostly EGFRvIII in G6^32^) and pT693-EGFR showed a downward trend without statistical significance in the disseminated tumor cells relative to primary tumor cells by INSIGHT, flow cytometry, and co-IF (**Fig. 4i**, **Extended Data Fig. 5a-f**). Notably, G6 disseminated tumor cells migrating as singlets exhibited lower, but detectable, EGFR and pT693-EGFR expression compared to those traveling in clusters, suggesting that the EGFR pathway is downregulated when cells disperse (**Extended Data Fig. 5c-f**). The downregulation of EGFRvIII expression in primary GBM at recurrence has been reported in patients^41–43^, drawing parallels between G6 disseminated tumors and the recurrent patient tumors originating from infiltrating cells post-surgery. Reductions in EGFR and pT693-EGFR expressions were also seen in G12 disseminated tumor cells compared to G12 primary tumor cells using microscopy, flow cytometry, and INSIGHT (**Extended Data Fig. 4e**, **Extended Data Fig. 6a-d**). Furthermore, disseminated G12 tumor cells, unlike G6, showed reduced Lamin A/C expression compared to primary cells by both INSIGHT and co-IF, while housekeeping proteins remained stable (**Extended Data Fig. 4e-f**). These findings reveal differences between disseminated G6 and G12 cells. EGFR downregulation was reported to facilitate epithelial to amoeboid transition during tumor invasion^44^, suggesting that EGFR signaling downregulation may be part of the G6 and G12 dissemination process regardless of EGFR mutation status. Results across multiple modalities revealed EGFR signaling suppression in disseminating GBM cells, confirming the robustness of INSIGHT and suggesting that disseminated PDX cells partially resemble recurrent patient GBM.

### Disseminated GBM cells increase postsynapse-associated program

Next, to uncover signaling circuitries that are altered with tumor spread, we compared the signaling networks of primary and disseminated tumor cells. G6 primary and disseminated tumor cells showed phosphorylation and protein expression profiles with subtle differences, suggesting disseminated cells largely retained the proteomic profile of primary tumor cells (**Extended Data Fig. 7a**). However, when we assessed the subtle but coordinated shifts in protein expression using by Gene Set Enrichment Analysis (GSEA), we saw the enrichment of pathways related to neuronal programs, including postsynaptic signal transmission, neurotransmitter receptors, and potassium channels, in disseminated cells (**Fig. 5a**). These findings align with studies showing that GBM cells can form functional synapses with neurons to promote tumor progression, affecting patient outcome^11,17–19,22^. Disseminated G12 cells exhibited more dynamic changes in protein expression compared to those of G6 models (**Extended Data Fig. 7b**), yet also showed enrichment of similar GO terms related to neuronal function and postsynaptic transmission in disseminated tumor cells relative to primary tumor cells (**Extended Data Fig. 7c**). Biological process involving regulation of postsynaptic membrane potential was enriched in disseminated G6 and G12 tumor cells compared to their primary counterparts, indicating enhanced postsynaptic regulation irrespective of EGFR mutation status (**Fig. 5b**). Top-ranked genes contributing to the enriched postsynaptic signatures in G6 disseminated cells were linked to EPH-ephrin-mediated repulsion and ion homeostasis, which have been implicated in tumor invasiveness^45–47^(**Extended Data Fig. 7d, e**). This prompted us to examine the synaptic genes in signatures related to tumor spread, including migration, invasion, and epithelial-to-mesenchymal transition. Querying the MSigDB database using keywords, we found that 45% of synaptic genes overlapped with the genes in tumor dissemination-related signatures (**Extended Data Fig. 8a**). Moreover, some phosphoproteins (e.g., MAPK1, DLG2) differentially expressed in disseminated cells compared to primary tumors appeared in multiple signatures (**Extended Data Fig. 8b**). These findings highlight an unrecognized extent of crosstalk between signaling circuitries governing tumor invasiveness and neuronal programs.

**Figure 5.**
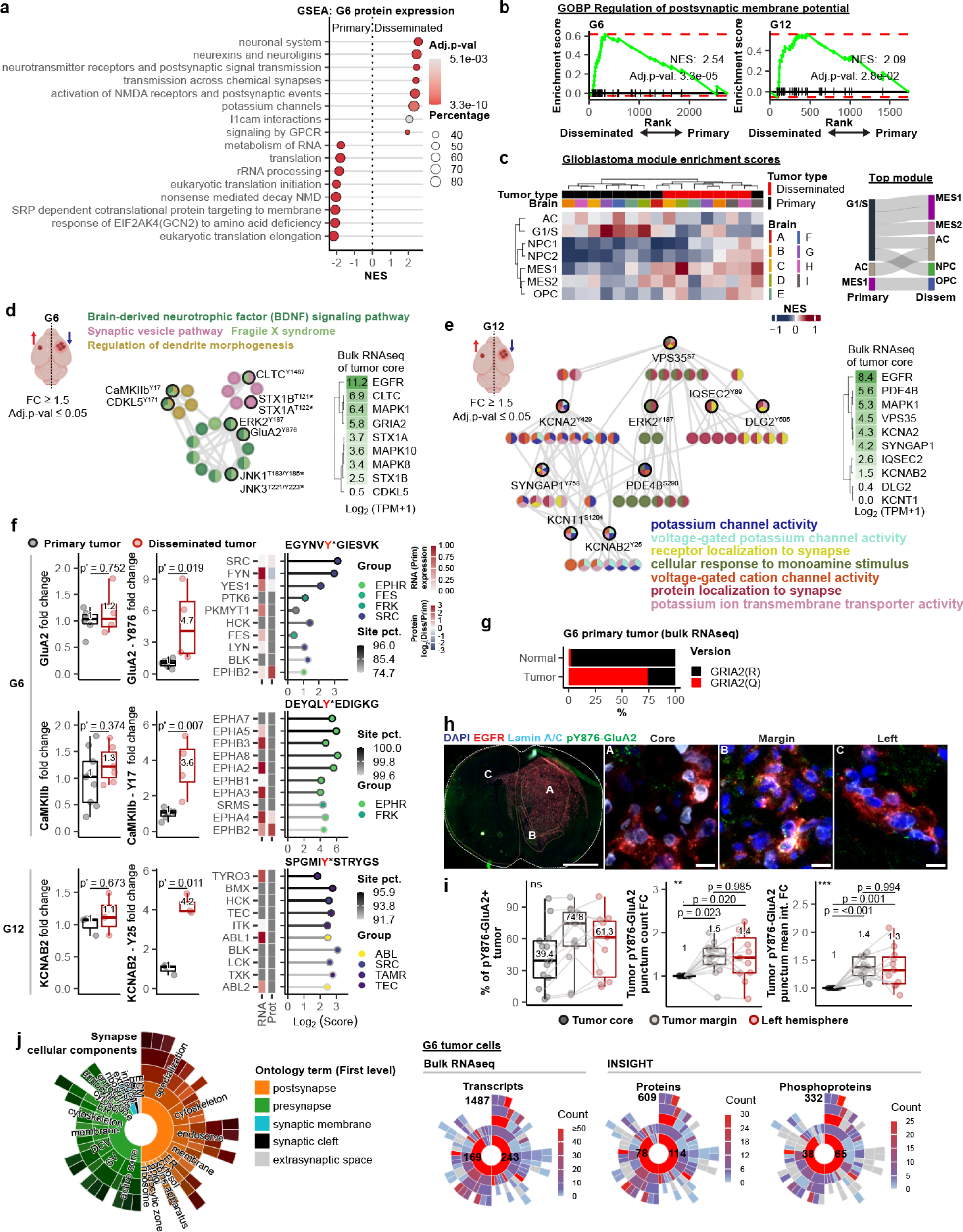
Disseminated GBM cells engage in postsynapse-associated signaling *in vivo*. (a) Gene Set Enrichment Analysis (GSEA) results on the proteins quantified in the primary and disseminated G6 tumors (log_2_ protein expression ratios) based on the Reactome database. (b) GSEA plots for the gene set “GOBP Regulation of postsynaptic membrane potential” comparing disseminated versus primary tumor cells. Enrichment scores are shown on the y-axis, with the ranked list of genes from disseminated (left) to primary (right) on the x-axis. The green line indicates the running enrichment score, while the vertical black lines represent individual genes contributing to the enrichment. The normalized enrichment score (NES) and adjusted p-value (Adj. p-val) for each PDX model are indicated. (c) Glioblastoma module^48^ enrichment scores (normalized enrichment score; NES) of primary and disseminated G6 tumor populations based on their protein expression as quantified by INSIGHT. Proteins belonging to NPC (neural progenitor cell-like), OPC (oligodendrocyte progenitor-like), MES (mesenchymal cell-like), G1/S, and AC (astrocyte-like) signatures were used. The Sankey plot (right) shows the state transition between the matching pairs of primary and disseminated tumor cells (n=7) based on their top enriched module. (d) Pathway enrichment analysis of phosphoproteins significantly upregulated (fold-change ≥ 1.5, adjusted p-val ≤ 0.05) in disseminated tumors compared to primary tumors in the G6 PDX model. Nodes represent phosphoproteins, with colors corresponding to pathway memberships. Phosphorylation sites are indicated for each protein. Pathway terms with the lowest p-values were annotated, and those that shared the phosphoproteins and had similar pathway terms are shown as extra nodes. *Due to shared tryptic phosphopeptide sequences between JNK1 and JNK3 as well as STX1A and STX1B, their protein identities could not be distinguished. Heatmap on the right shows the expression level (Log_2_(TPM+1)) of the corresponding human-specific transcripts (and EGFR as reference) from bulk RNAseq of core tumor tissue^30^. TPM = Transcripts Per Million. (e) Pathway enrichment analysis of phosphoproteins significantly upregulated (fold-change ≥ 1.5, adjusted p-val ≤ 0.05) in disseminated tumors compared to primary tumors in the G12 PDX model as done in **d**. (f) Box plots (left) of select differentially expressed phosphorylation sites and their corresponding protein level in disseminated (red) and primary (gray) tumor samples of G6 (pY876-GluA2, pY17-CaMKIIb) and G12 (pY25-KCNAB2) PDX models. BH adjusted p-values (p’) are shown. The lollipop plots (right) represent candidate kinases for the matching phosphorylation sites (left) based on the Kinase Library prediction. The phosphopeptide queried is shown at the top of the lollipop plots, with the phosphorylated residue indicated in red with asterisk. The site percentile (pct.) indicates the percentile score of the peptide in the phosphoproteome as the substrate for the kinase listed. Heatmaps left to the lollipop plot indicate the scaled level of transcript (RNA) in the tumor core from bulk RNAseq, and relative log_2_(disseminated/primary tumor) values (Prot) from protein expression data for each kinase. Gray cells in heatmap indicate proteins undetected by INSIGHT or transcripts with Log₂(TPM + 1) < 0.1. TPM = Transcripts Per Million. (g) Percentages of GluA2(Q) and GluA2(R) in primary G6 tumor tissue based on all human *GRIA2* reads by bulk RNAseq. (h) Co-IF of the whole coronal section (first image) of G6 PDX-tumor bearing brain. Regions of interest labeled: A (tumor core), B (tumor margin), C (corpus callosum in the left hemisphere). Staining included DAPI (blue, nuclei), human EGFR (red), human Lamin A/C (cyan), and pY876-GluA2 (green). Representative images are shown. Scale bar = 10 µm for magnified images and 2mm for whole section. Continued in **Extended Data Figure 9d, e**. (i) Quantification of the co-IF analysis done in **h**. (Left) Percentage of pY876-GluA2 positive tumor cells in the tumor core (black), tumor margin (gray), and left hemisphere (red). (Middle) Fold change (FC) in pY876-GluA2+ punctum counts on tumors across tumor core, margin, and left hemisphere. (Right) Mean fluorescence intensity fold change of pY876-GluA2 puncta on tumors across regions. ANOVA p-value (*** = p-value ≤ 0.001; ** = p-value ≤ 0.01; ns = not significant) is shown at the top left. Adjusted p-values between conditions shown. (j) Ontology analysis of synaptic components (annotated in SynGO^69^) in G6 primary and disseminated tumor cells using bulk RNAseq and INSIGHT datasets. The left sunburst plot illustrates the first-level ontology terms related to synaptic structures (cellular components). The right panels show the count distribution of transcripts (bulk RNAseq), proteins, and phosphoproteins (INSIGHT) annotated with each ontology term, using concentric heatmaps. The red-to-blue color scale represents the count per term, with central numbers indicating counts for presynapses and postsynapse GO terms. The numbers at the top represent all the synapse-associated terms found in the data.

Cell state shifts between disseminated GBM cells and primary GBM cells at the protein level *in vivo* have remained unclear. The enrichment analysis of GBM cell state modules^48^ based on the proteome of tumor cells revealed a shift from G1/S or G2/M cell cycling states to mesenchymal or neuronal progenitor-like cell states in G6 and G12 models upon dissemination (**Fig. 5c, Extended Data Fig. 7f**). These results align with studies showing that recurrent GBM acquires mesenchymal cell state^49,50^ while invasive GBM cells acquire neuronal progenitor-like cell state^22^ based on transcriptome data. These findings demonstrate that cell state modules based on proteins and transcripts align and that disseminated GBM cells resemble invasive or recurrent GBM.

### Phosphorylation-driven modulation of synapse-associated signaling underpins GBM dissemination

In the G6 model, phosphorylation of proteins linked to synapse function and neuronal programs was significantly increased in disseminated tumors compared to primary tumor cells (**Fig. 5d**). Some of these synapse-associated proteins also had high gene expression in primary tumor samples *in vivo*, based on their relative transcript levels to that of amplified EGFR, reinforcing that primary tumor cells can express them (**Fig. 5d**). Disseminated G6 tumors also showed a subtle increase in the phosphorylation of top-ranking GSEA-enriched postsynaptic proteins relative to primary G6 tumors (**Extended Data Fig. 7e right**), together indicating that tumor cells engage postsynapse-associated signaling networks as they spread. Disseminated G12 tumor cells similarly increased postsynapse-related signaling that is centered around ion homeostasis relative to its primary counterpart (**Fig. 5e, Extended Data Fig. 9a**). Notably, in both models, although the expressions of many total proteins were comparable between primary and disseminated tumor cells, the phosphorylation of various postsynaptic proteins including pY876-GluA2, pY17-CaMKIIb, pY171-CDKL5, pY25-KCNAB2, pY758-SYNGAP1 was elevated (**Fig. 5f, Extended Data Fig. 9a-c**), underscoring that disseminated GBM cells co-opt synapse-associated programs that are regulated by phosphorylation rather than protein expression, which have not been previously recognized. Proteins that regulate neuronal migration, including SYNGAP1, CLTC, CDKL5, and CaMKIIb, were highly phosphorylated, suggesting that invasive tumor cells likely co-opt neuronal migratory signaling pathways, strongly aligning with a recent study^22^, and implicating those phosphorylation sites in GBM dissemination for the first time. Intriguingly, in G6, total protein and doubly phosphorylated T185/Y187-ERK2 (undetected in G12) remained unchanged between primary and disseminated tumors while pY187-ERK2 increased, indicating selective and differential phosphorylation on ERK2 during tumor spread, a phenomenon that has not yet been explored (**Extended Data Fig. 9c**). Kinase Library^4,5^ predicted that Src phosphorylates Y876-GluA2 (**Fig. 5f**), consistent with previous reports^51–53^. Src was detected by both RNAseq (primary tumor) and INSIGHT in G6 primary and disseminated tumors (**Fig. 5f**), supporting the potential role of Src in GluA2 phosphorylation *in vivo*. EPHR kinases were predicted to potentially phosphorylate Y17-CaMKIIb; accordingly, EPHB2 was elevated (without statistical significance) in disseminated tumor cells relative to primary tumor cells (**Fig. 5f**). Additionally, phosphorylation of KCNAB2, a potassium channel recently implicated GBM invasion^54^ and elevated in G12 disseminated tumor cells, ranked highly as a substrate of TYRO3, a kinase involved in synaptic plasticity and tumor invasion^55,56^. This places KCNAB2 and TYRO3 at the cross-section of the two signaling circuitries and implicates pY25 in GBM dissemination (**Fig. 5f**). Together, our findings uncover numerous previously uncharacterized phosphorylation sites during GBM dissemination and collectively highlight that phosphorylation-driven modulation of synapse-associated signaling circuitries underpins GBM dissemination.

### Disseminated GBM cells increase Y876-GluA2 phosphorylation

The phosphorylation of Y876-GluA2 is essential for synaptic plasticity in neurons^57–61^, but its role in cancer is unknown. Thus, we set out to validate that it is indeed upregulated in disseminated GBM cells. Normally, RNA editing of GluA2 converts its glutamine (Q) to arginine (R), rendering it Ca^2+^-impermeable^62^; however, in glioma, reduced editing increases GluA2(Q) expression^17,62^, which has been shown to enhance Ca^2+^ influx and allow synaptic transmission-dependent tumor growth and synaptic transmission-independent invasion and migration^63,64^. Previous studies showed that GluA2 is absent in GBM cell lines^65^ but present in primary human GBM^17,63^, indicating that G6 PDX tumor cells capture a feature of human GBM *in vivo*. A bulk RNAseq analysis of primary G6 tumor cells revealed that GluA2(Q) transcripts constitute a striking 74% of the total *GRIA2* expressed, indicating a predominant expression of Ca^2+^-permeable GluA2 by G6 tumor cells *in vivo* (**Fig. 5g**). By co-IF, the percentage of tumor cells that contained at least one pY876-GluA2 positive puncta did not significantly differ across regions (**Fig. 5h, i, Extended Data Fig. 9d, e**). However, consistent with the INSIGHT data, the number and the mean intensity of pY876-GluA2-positive puncta were significantly higher in tumor cells at the margin and the left hemisphere compared to those at the core, suggesting tumor cells may leverage pY876-GluA2 to regulate synaptic plasticity as they spread (**Fig. 5h, i, Extended Data Fig. 9d, e**). Together, our findings show that GBM cells highly express Ca^2+^-permeable GluA2 and can increase pY876-GluA2 at the tumor margin and maintain it in the left hemisphere, implicating pY876-GluA2 in GBM dissemination.

### GBM cells express both presynaptic and postsynaptic proteins but differentially phosphorylate postsynaptic proteins during dissemination

Although GBM has been shown to express both presynaptic and postsynaptic proteins^66–68^, their general function and whether they are differentially regulated during GBM progression remain largely unknown. As most of the significantly upregulated synaptic phosphoproteins in disseminated tumor cells relative to primary tumor cells were associated with postsynapse function, we examined the synapse component memberships of proteins and phosphoproteins expressed in G6 and G12 tumors using SynGO^69^. Both bulk RNAseq and INSIGHT quantified a broad spectrum of both presynapse and postsynapse-associated proteins and phosphoproteins in both G6 and G12 primary and disseminated tumors (**Fig. 5j**, **Extended Data Fig. 9f**). These findings indicate that, while GBM cells indeed express both presynaptic and postsynaptic proteins and phosphoproteins, postsynaptic proteins exhibit distinct phosphorylation dynamics during dissemination, underscoring their role in regulating GBM dissemination.

### Cell cycle program is downregulated in disseminated GBM cells

Next, to understand which signaling circuitries are suppressed with tumor spread, we analyzed proteins with reduced phosphorylation in disseminated tumor cells relative to primary tumor cells. We found highly interconnected circuitries associated with proliferation in both models (**Fig. 6a, Extended Data Fig. 10a**). Moreover, MYC transcriptional targets, crucial for cell proliferation^70^, were downregulated in disseminated cells of both models compared to their primary counterparts by GSEA (**Fig. 6b**). These findings aligned with the GBM module enrichment result where G6 and G12 primary tumor cells shifted from G1/S or G2/M cell states to other cell states upon dissemination (**Fig. 5c, Extended Data Fig. 7h**). Notably, disseminated G6 tumor cells had reduced pT373-RB1 expression (undetected in G12) (**Fig. 6c**). RB1 phosphorylation at T373 by CDKs allows G1/S phase cell cycle progression and cell proliferation^71–73^. Indeed, by co-IF, the percentage of nuclear pT373-RB1-positive tumor cells was significantly decreased in the disseminated tumors relative to primary tumors in both G6 and G12 models (**Fig. 6d-g, Extended Data Fig. 10b**). Probing for pS807/811-RB1, which follows pT373-RB1 to secure G0 phase exit, similarly showed a marked decrease in disseminated tumor cells in G6 and G12 relative to their primary tumors^74,75^ (**Fig. 6d-g, Extended Data Fig. 10c**). Thus, our findings indicate that disseminating tumor cells undergo reduced cell cycling. These results support reports that tumor cells shift between proliferative and migratory states, decreasing proliferation to invade surrounding tissues^76,77^, and highlight that this transition is also reflected in their signaling networks at the systems level *in vivo,* which has not been previously recognized. Interestingly, we observed a smaller reduction of percent positive pS807/811-RB1, along with greater sustainment of pS807/8011 and pT373 levels in G6 compared to G12, indicating that G6 tumor cells likely continue to proliferate in the left hemisphere after dissemination (**Fig. 6d-g**). This aligns with the greater dissemination observed in G6 compared to G12 (**Fig. 3a, Extended Data Fig. 3a-b**). In addition, similar to pY876-GluA2 (**Fig. 5h, i**), pRB1 in tumor cells was reduced at the margin (**Fig. 6e, g**), indicating that tumor cells at the margin resemble those in the left hemisphere.

**Figure 6.**
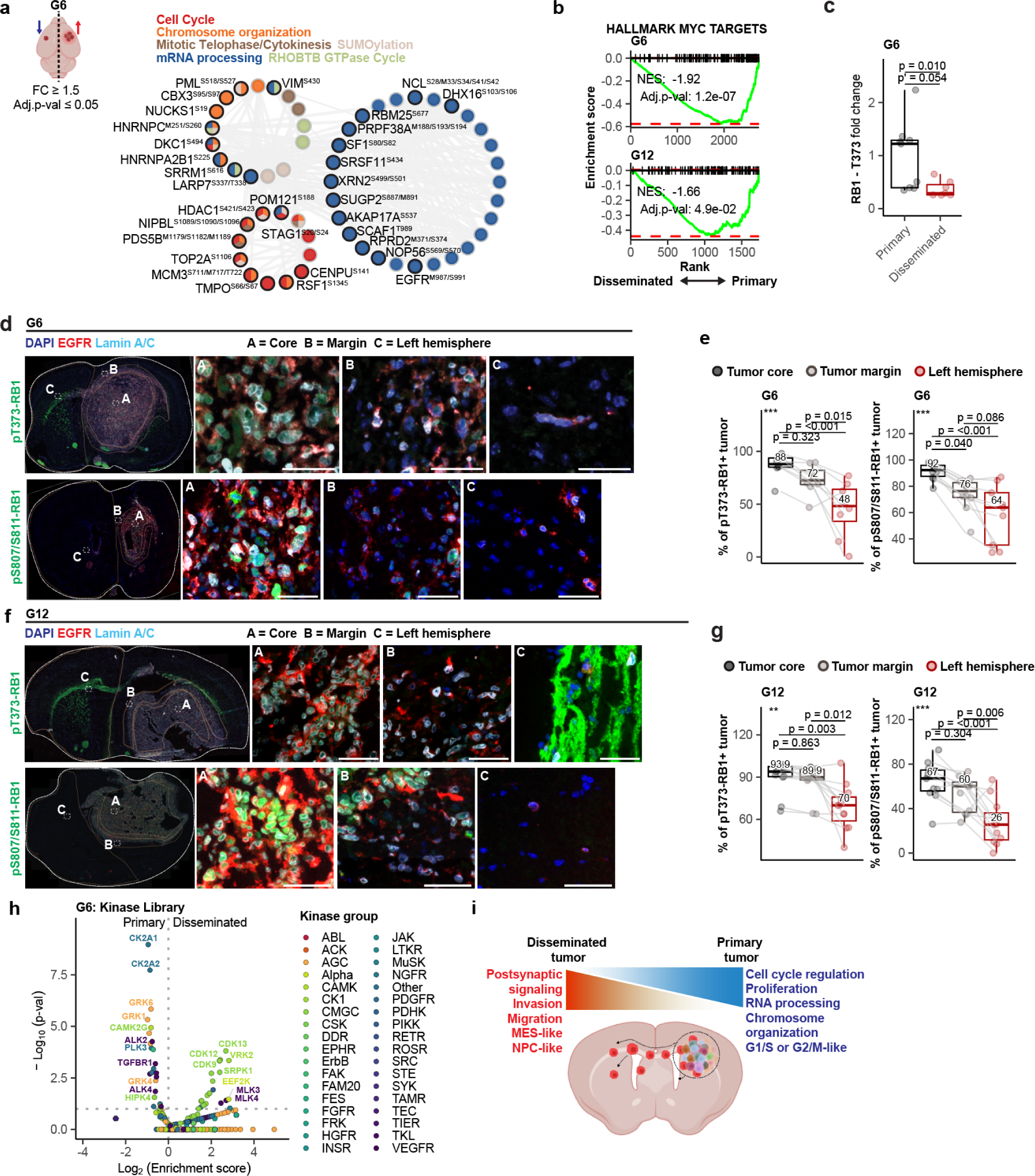
Proliferation-associated signaling network is suppressed in disseminated GBM cells. (a) Pathway enrichment analysis of phosphoproteins significantly downregulated (fold-change ≥ 1.5, adjusted p-val ≤ 0.05) in disseminated G6 tumors compared to primary G6 tumors. Membership of the phosphoproteins (nodes) in each pathway is denoted by the same colors as the pathway annotated (top). Phosphorylation sites are indicated in superscript. Pathway terms with the lowest p-values were annotated, and those that shared the phosphoproteins and had similar pathway terms were shown as extra nodes. (b) GSEA plots for the gene set “Hallmark Myc targets v1” comparing disseminated versus primary tumor cells. Enrichment scores are shown on the y-axis, with the ranked list of genes from disseminated (left) to primary (right) on the x-axis. The green line indicates the running enrichment score, while the vertical black lines represent individual genes contributing to the enrichment. The normalized enrichment score (NES) and adjusted p-value (Adj. p-val) for each sample (G6 and G12) are indicated. (c) Box plot shows the fold changes of pT373-RB1 as quantified by INSIGHT of G6 tumor cells. BH-adjusted p-values (p’) and unadjusted p-value (p) are shown. (d) G6 PDX tumor-bearing brains analyzed by co-IF. Staining for pT373-RB1 (green; top), and pS807/S811-RB1 (green; bottom), with DAPI (blue, nuclei), human EGFR (red), and human Lamin A/C (cyan) shown. The whole section (left) with dotted lines delineates the left hemisphere, the core, and the invasive margin of the primary tumor. The magnified regions (right) shown are labeled in the whole section: A (tumor core), B (tumor margin), and C (left hemisphere). Scale bar = 40 µm. Continued in **Extended Data Figure 10b, c**. (e) The percentage of pT373-RB1+ and pS807/811-RB1+ G6 tumor cells in the tumor core, invasive margin, and the left hemisphere stained in d. ANOVA p-value (*** = p-value ≤ 0.001; ** = p-value ≤ 0.01) is shown at the top left corner. Adjusted p-values between conditions shown. (f) Same analysis as in panel **d** was performed for G12 PDX tumor-bearing brains. Continued in **Extended Data Figure 10b, c.** (g) Same analysis as in panel **e** was performed for G12 PDX tumor-bearing brains stained in **f**. (h) Kinase library analysis of differentially expressed phosphoproteome between primary and disseminated G6 tumor cells (1.5-fold change, adj. p-val ≤ 0.1) to infer enriched active S/T and Y kinases. X-axis represents enrichment scores calculated by Kinase Library (i) Disseminated (left hemisphere) and primary tumors (right hemisphere) as characterized by INSIGHT on GBM PDX models. Tumor cells (left, red) increasingly engage in postsynapse-associated signaling circuitries as they disseminate the normal brain, while primary tumor cells (right, blue) engage in proliferation-associated signaling circuitries.

In both PDX models, hornerin (HRNR)—a calcium-binding S100 protein implicated in cell cycle regulation^78^ and invasive tumor progression^79–83^—exhibited the highest upregulation during dissemination. This suggests HRNR, previously unassociated with GBM, may contribute to the invasive phenotype of disseminated GBM cells *in vivo* (**Extended Data Fig. 10d**). CDK5, a regulator of neuronal migration and synaptic plasticity^84^, was predicated as a kinase for HRNR phosphorylation, suggesting a potential cross-talk between neuronal, cell cycle, and invasion-associated signaling circuitries (**Extended Data Fig. 10e**).

### Primary and disseminated glioblastoma cells are predicted to have differential kinome activity

Global changes in kinase activities between disseminated and primary GBM *in vivo* remain unknown despite their potential therapeutic implication. The inference of differentially active kinases between disseminated and primary G6 tumor cells based on their phosphoproteomes revealed VRK2 and MLK4 as some of the top-ranking kinases in disseminated tumor cells (**Fig. 6h**). Both are preferentially enriched in stem-like mesenchymal GBM tumors^85^, with VRK2 linked to glioma cell cycle regulation^86^ and MLK4 being essential for mesenchymal GBM survival^85^, raising the possibility that other inferred kinases with unknown functions in GBM may also contribute to dissemination.

## Discussion

We developed INSIGHT, a platform technology that enables discovery-based, systems-level signaling network characterization of specific cell types or subpopulations from complex tissues without requiring genetic engineering, positioning it as a valuable tool for both basic and translational research. The consistency between orthogonal experimental methods across key findings on GBM PDX models highlights the robustness of INSIGHT even for rare cell subsets. INSIGHT also offers opportunities for monitoring perturbations of kinases or phosphatases and their phosphorylated substrates specific to those cells, which could enhance our understanding of heterogeneous cellular functions and precision in therapeutic targeting *in vivo*. Furthermore, inferring ligand-receptor interactions from signaling networks as we have done will enhance our understanding of cell-cell communication, as they directly reflect the biochemical states of cells. The development of computational tools that optimally leverage such data will be important. Moreover, INSIGHT overcomes various challenges faced by bulk phosphoproteomics of PDX models^35^, including distinguishing human and mouse proteins, highlighting its utility in phosphoproteomics of mixed-species models.

Using INSIGHT on preclinical GBM PDX models, we characterized thousands of phosphoproteins and proteins within disseminated GBM, providing new insights into the signaling network underpinning GBM dissemination. Systems-level analysis of GBM signaling network rewiring *in vivo* with tumor spread showed a global transition from proliferative to postsynapse-associated signaling (**Fig. 6j**). We also highlight that signaling circuitries of neuronal and non-neuronal programs interconnect during GBM dissemination by uncovering numerous phosphoproteins, proteins, and kinases and their relationships within those circuitries. Alterations in phosphorylation rather than total protein expression were seen in many signaling components with rewiring, underscoring the importance of quantifying post-translational modifications *in vivo* to fully understand disease progression. Importantly, our work points to a previously unrecognized role of pY876-GluA2 in GBM dissemination. In neurons, pY876-GluA2 mediates synaptic upscaling and long-term potentiation (LTP) by promoting the accumulation of GluA2 at the postsynaptic density^57–61,87,88^. Phosphorylation of Y876-GluA2(Q) may regulate neuron-GBM synaptic plasticity through similar mechanisms, particularly for those GBM cells with multisynaptic contact with neurons^19^, facilitating tumor growth or invasion via neuronal interaction, while separately promoting invasion through neuron-independent but GluA2(Q)-dependent mechanisms that rely on increased GluA2 at the membrane. Further studies are needed to ascertain whether phosphorylation occurs on both GluA2(Q) and GluA2(R) and to define their functional relevance in GBM dissemination. Additionally, the relevance of interrogating the systems-level signaling networks within contralaterally disseminated GBM PDX cells for insights into invasive and recurrent GBM biology has been unclear. We clarify this by demonstrating that key phosphoproteins show consistent regulatory patterns in both disseminated cells and those at the tumor margin—the dominant site of recurrence^14^. Disseminated cells also adopt a mesenchymal cell state and downregulate EGFR signaling, reflecting features of recurrent GBM^41–43,49,50^. Combining INSIGHT with microscopy-based validation therefore may offer a framework for investigating invasive GBM mechanisms contributing to recurrence.

Refinement of the INSIGHT will increase its utility. First, fixation can negatively impact the epitope recognition of some antibodies. Using antibodies targeting cytoplasmic markers and those used for immunohistochemistry will broaden the range of detectable markers. Second, we employed data-dependent acquisition, which, while effective for discovery-based proteomics, is inherently limited by stochastic precursor selection, impacting the reproducibility across experiments. Targeted acquisition strategies will increase sensitivity, reproducibility, and precision^89,90^. Third, dyes that rely on membrane integrity for live/dead cell discrimination become ineffective with fixation. While INSIGHT performed effectively in both normal and tumor-bearing tissues, strategies to discern live/dead cells without relying on membrane impermeability would minimize noise. Fourth, given that INSIGHT utilizes cryosections, sampling parallel tissue sections with spatial-omics technologies will increase the depth of biological insight. Even with these current considerations, INSIGHT advances *in vivo* MS-based phosphoproteomics for targeted, cell subset-specific signaling network analysis, providing a platform to uncover mechanistic insights with broad applications in both basic and translational research.

## Methods

### Animal studies

GBM6 (G6) and GBM12 (G12) PDX tumor intracranial inoculations were performed in female Nude mice as previously described^91,92^. G6 was maintained in FBS-based media, and G12 was maintained in stem cell media. TdTomato-positive G6 PDX tumor cells were generated as previously described^93^. All animal work related to PDX-tumor generation was done in compliance with the Mayo Institutional Animal Care and Use Committee. Detailed information on the clinical and molecular characteristics of G6 and G12 PDXs is in **Supplementary Table 1** (summarized from the information available on the Mayo Clinic Brain Tumor PDX National Resource maintained by the Sarkaria lab). For the studies using normal organs, male and female C57BL/6 mice were acquired from Taconic Biosciences, maintained, and necropsied in compliance with protocols approved by the Committee on Animal Care (CAC/IACUC) at MIT.

### Isolation of single cells from fixed tissue

Flash-frozen brains (normal or PDX-tumor bearing) embedded in OCT compound were cut down the mid-sagittal plane, and each hemisphere was sectioned using a Cryostat (Leica) and fixed. Normal brain and PDX tumor-bearing brain tissues were washed after fixation and incubated with collagenases. Undissociated tissues were briefly sonicated in specific locations within a bath sonicator (VWR) to yield single-cell suspension.

### Co-immunofluorescence microscopy and analysis

Frozen whole brains embedded in OCT compound were coronally sectioned at 5 µm, and Lamin A/C (Abcam) stain was performed at intervals to identify the location of the most prominent primary tumor and disseminated tumor to count the number and percentage of tumor cells in each region identified. Two sections from two different coronal depths were stained and analyzed per PDX tumor-bearing brain for 6 brains (total n = 12 sections). Tumor burden varied across depths. When disseminated tumor cells were absent, analysis of the primary tumor and margins was still performed. Sections lacking a clear midline to delineate the left hemisphere or those with extensive primary tumor growth that pushed into the left hemisphere were excluded, consistent with the INSIGHT analysis. The tumor core, margin, and left hemisphere were drawn manually for each section for analysis. Tissues were fixed with 2% paraformaldehyde/PBS, washed with PBS, permeabilized with 0.1% Triton100/PBS, and blocked with normal serum. Tissues were stained with primary antibodies targeting the pT373-RB1, pS807/811-RB1, pY876-GluA2, pT693-EGFR and were probed with goat anti-rabbit IgG-AF488 (H+L) secondary antibody. We note that strong staining in the corpus callosum was seen for the Abcam pT373-RB1 antibody. Sections were blocked with normal serum and incubated with rabbit anti-human Lamin A/C-AF594 and mouse anti-human EGFR-APC. Sections were stained with DAPI after being subjected to TrueView (Vectors) to distinguish nuclei, mounted using Vectashield vibrance antifade mounting medium, and images were taken using a TissueFAXS SL slide scanner (TissueGnostics) coupled to a Zeiss upright microscope with a 20x or 40x objective with z-stack enabled. All images from TissueFAXS SL were analyzed using StrataQuest (v7.1.1.143). A nuclei mask was used for the quantification of pT373-RB1 and pS807/811-RB1 and Lamin A/C while a cellular mask was used for the quantification of EGFR and pT693-EGFR. For pY876-GluA2, the FISH dot algorithm was used. For all image analyses of disseminated tumors in the left hemisphere of the G12 PDX model, each tumor cell was visually validated. For confocal microscopy, Olympus FV1200 Laser Scanning Confocal Microscope was used at 60x objective with 2x zoom, and the images were analyzed using Fiji.

### Liquid chromatography-tandem mass spectrometry (LC-MS/MS)

Samples were lysed in 50mM Tris/5%SDS, heated, sonicated, reduced with 10mM DTT, alkylated with 55mM IAA, and bound to s-trap (Protifi), washed with 50% chloroform/50% methanol, washed with 100mM TEAB/90% methanol, and digested with trypsin overnight before elution. For TMT labeling, lyophilized peptides were resuspended in 50mM HEPES and labeled with TMT (Thermo). For tyrosine phosphoproteome, the enrichment of pY peptides was performed by resuspending the TMT labeled samples in immunoprecipitation buffer (Tris-HCl/1% NP-40) and incubating digested peptides with protein G agarose beads pre-conjugated to 4G10 and PT66 antibodies overnight. Subsequently, immunoprecipitated peptides were eluted with trifluoroacetic acid and enriched for phosphopeptides using ferric nitrilotriacetate columns (Thermo). Eluates were dried, reconstituted, and manually loaded using a column-packing device onto a 12 cm analytical column containing C18 beads freshly hand-packed and conditioned on the day of each experiment. Analytical columns were laser-pulled, fritted with Lithisil and formamide, and flamed to open the frits on the day of each experiment. Agilent 1100 Series HPLC connected to Orbitrap Exploris 480 Mass Spectrometer was operated at 0.2 ml/min flow rates with a precolumn split to attain nanoliter flow rates through the analytical column and nano-electrospray ionization tip. Peptides were eluted with the increasing concentrations of buffer B using the following 140-minute gradient settings (Buffer A: 0.1% acetic acid; Buffer B: 70% acetonitrile/0.1% acetic acid): 0 min: 0% B; 10 min: 11% B; 110 min: 32% B; 125 min: 60% B; 130 min: 100% B; 128 min: 100% B; 130 min: 0% B. The MS parameters were: ESI spray voltage, 2.5 kV; no sheath or auxiliary gas flow; heated capillary temperature, 275 °C, data-dependent acquisition mode with the scan range of 380–2000 m/z. MS1 scans were acquired at 60,000 resolution, maximum injection time of 100 ms, normalized AGC target of 300%, and only included the precursor charge states of ≥ 2 and ≤ 6. For every full scan, MS/MS spectra were collected during a 3-second cycle time. For MS/MS, ions were isolated (0.4 m/z isolation width) with the maximum IT of 250 ms, normalized AGC target of 100%, HCD collision energy of 33% at a resolution of 60,000, and the dynamic exclusion time of 35 seconds. Half of the supernatant from pY-IP was subjected to high-pH reverse-phase fractionation on a Kromasil® C18 HPLC Column (5 μm particle size, pore size 100 Å, L × ID 250 mm × 4.6 mm) using buffer A (10 mM triethylammonium bicarbonate (TEAB), pH 8) and buffer B (10 mM TEAB, pH 8, 99% acetonitrile) over an 85-minute gradient (0 min: 1% B, 1 min: 1% B, 5 min: 5% B, 65 min: 40% B, 75 min: 70% B, 84 min: 70% B, 85 min: 1%) into 10 fractions (concatenated) using a Gilson FC 204 Fraction Collector and Agilent 1100 Series HPLC. Fractionation was performed at a flow rate of 1ml/min and 1 min per fraction between 10-85min portion of the gradient. For each fraction, 1/10 was allocated for protein expression analysis, and 9/10 was allocated for global phosphoproteomic (pS/pT and residual pY not already captured by 4G10 and PT66 antibodies) analysis. For simplicity, data generated from the global phosphoproteomic analysis were denoted as “pS/pT” while those from pY-IP-enrichment were denoted as “pY” in the study. Fractions for global phosphoproteome analysis were resuspended in 0.2% TFA and subjected to Fe-NTA column-based phosphopeptide enrichment, after which they were eluted, dried, and resuspended for injection by Dionex UltiMate 3000 Autosampler (Thermo) on a 1.9 μm C18 bead-packed analytical columns.

### Mass spectrometry data analysis

Mass spectra were searched using Mascot (v2.8) and Proteome Discoverer (v3.0) against the Swiss-Prot database (Mus musculus proteome for murine tissues, and concatenated Homo sapiens and Mus musculus proteome (2023_06 and 2024_03) appended with common Repository of Adventitious Proteins (cRAP) fasta sequences for PDX or mixed tissues). Raw files were searched with two or fewer missed cleavages, precursor and fragment ion matched with 10 ppm and 20mmu mass tolerances, and fixed modifications of carbamidomethyl (Cys), TMT-pro (+304.2071) at peptide N-terminus and lysine and dynamic modification of oxidation (Met). For phosphoproteomic data, additional dynamic modifications of phosphorylation (Ser, Thr, Tyr) were added. TMT reporter ion intensities were corrected with isotope correction factors provided by Thermo Fisher (lot # for each run can be found on PRIDE). A target-decoy search strategy was used to adjust Peptide Spectral Matches (PSMs) to 1% FDR, and the Percolator^94^ node was used in the processing step to improve the sensitivity and accuracy of peptide identification and the resulting q-values were used to keep only those PSMs with q-value ≤ 0.01 (“1% FDR” reported in figures). The ptmRS^95^ node was used to confidently assign the phosphorylated and oxidized sites (filtered for peptides with localization probability ≥ 90% after search). PSMs were further filtered with ion scores ≥20, a cutoff determined in-house through extensive manual validation of spectra for higher-stringency identification (“HS”) compared to 1% FDR^26^. TMT intensities for common PSMs were summed for peptide identification and quantification. Proteins mapped to only one protein accession per species were used for all analyses. For protein accessions without associated gene names, protein accession itself was used for graphing. The residue positions of the phosphorylation (Y, S, T) and oxidation (M) were extracted using the full-length sequence of the longest isoform of the protein. To adjust for differential sample loading, all protein expression and phosphoproteomic data were normalized first using the median intensity of each TMT channel from their matching protein expression data. To join all the multiplexed sets derived from the PDXs, the samples within each plex were normalized to the normal cells from the left hemisphere with the assumption that those cells stay relatively consistent across all brains. For the analysis of O4 positive cells, primary tumors, and normal cells from the left hemisphere, all missing values were filled in with the lowest TMT intensity from the corresponding spectrum that was extracted and divided 2-fold and used to impute the missing values (“MGF method”) as previously described^96^ with the assumption that the majority of these values were not detected because they were absent or low in abundance (e.g., O4-positive cells may not express proteins that O4-negative cells express, and vice versa). If the lowest m/z intensity was larger than the lowest isotope impurity-corrected TMT reporter intensity within the PSM, ½ of the latter was used instead. To do this, MGF (Mascot Generic Format) files were read in with the MSnbase (2.28.1)^97^, and the intensities were assigned based on the First Scan number. For the analysis of disseminated tumor cells, all PSMs with missing values across all samples were removed from the analysis (with the assumption that most of the missing value arose due to low sample input and not because they did not express those peptides), except for those PSMs where missing values existed only in samples other than the disseminated tumor samples (here the missing values were replaced using the MGF method as above). This was done to avoid bias introduced by imputation. This allowed the plotting of the relative changes of tumor-specific markers (human lamin A/C (LMNA), human OLIG2, human Nestin (NES), human EGFR) across all samples. To retain maximum information without imputing or excluding phosphopeptide or protein identifications (IDs) with missing values, IDs with equal combinations of sample sizes per condition were grouped and subjected to differential expression analysis. To remove potential doublets and debris that may be of murine origin for isolated tumor cells, the following criteria were used to filter the protein expression data for each multiplexed set: 1) when proteins arose from peptides that uniquely mapped to human (ph) or both human and mouse (phm) proteomes, only ph-derived proteins and their TMT intensities were kept; 2) when the same protein arose from both phm and pm, Pearson correlation analysis across all samples across plexes was done to remove the phm intensities that had a correlation coefficient (*r*) > 0 to pm regardless of the adjusted p-value, aside for those with 0 ≤ *r* ≤ 0.2 and adjusted p-value 0.3; 3) peptides that uniquely mapped to mouse proteome (pm) were removed. For the phosphoproteome data, all values associated with phosphopeptides whose total proteins in the protein expression data were filtered out using the aforementioned criteria were also removed. Furthermore, phosphopeptides that mapped to both human and mouse proteins were removed if the protein expression data only detected the murine version of its matching total protein. Phosphopeptides that were present in less than three disseminated tumor samples and three primary tumor samples were removed from consideration. EGFR-positive tumor populations from two left hemispheres (A and F) were excluded from data analysis, as they represented portions of the primary tumors that extended into the left hemisphere and inadvertently included during the mid-sagittal dissection, resulting in their high percentage of tumor cells compared to other left hemispheres during sorting (**Extended Data Fig. 3c**). The same phenomenon was observed in the batches of tumor-bearing brains analyzed by microscopy, and these cases were excluded from analysis as detailed in the microscopy methods section.

### Flow cytometry and sorting

All flow cytometric analyses were done using BD High Throughput Sampler (HTS) coupled to LSRFortessa (BD Biosciences). Anti-EGFR-PE antibody was used to stain and identify tumor cells from normal cells for the samples subjected to mass spectrometry. For all sorting experiments, Sony MA900 was used to sort cells. For testing fixatives, anti-EGFR-BV421 was used with a gating strategy shown in **Supplementary Figure 1a**. Mixture of normal and TdTomato+ tumor cells were stained with anti-HLA-A/B/C-APC-Cy7 antibody or anti-EGFR-BV421 antibody (gating strategy in **Supplementary Figure 1b**). For dissociated PDX tumor-bearing brains, cells were first gated on FSC-A vs. SSC-A to exclude debris and aggregates, followed by FSC-W vs. FSC-H and SSC-H vs. SSC-A to isolate singlets, after which cells were gated on EGFR-PE vs. SSC-A to separate EGFR-positive and EGFR-negative populations (**Supplementary Figure 1c**).

### RNAseq analysis

Previously published bulk RNAseq data^30^ for intracranially implanted core G6 and G12 PDX tumors were retrieved and re-processed. Data processing was done in part using Nextflow-based nf-core workflows^98^. RNA-seq data from bioproject PRJNA548556 with sample IDs SRR9294059 (G6) and SRR9294060 (G12) were obtained using the nf-core/fetchngs version 1.12.0^99^ and used to quantify transcripts from a combined human hg38 and mouse mm39 genome assembly with Ensembl version 112 annotations using the nf-core/rnaseq workflow revision 3.14.0^100^. Gene level summaries were prepared from the star_salmon quantitation using tximport version 1.32.0^101^ running under R version 4.4.1^102^ with tidyverse version 2.0.0^103^. Genomic alignments were visualized using the Integrated Genomics Viewer version 2.17.0^104^. Human-specific protein-coding gene expression values were reported as log_2_-transformed Transcripts Per Million (TPM) with a +1 offset in all figures and tables (**Supplementary Table 3**). For the analysis of amino acid 607 codon sequence coverage in human *GRIA2* and mouse *Gria2*, reported percentages included all aligned reads.

### Data analysis

All data were processed and analyzed using R (v4.3.1) and R studio. Differential expression analysis was performed using the limma R package^105^ with a significance cutoff of adjusted p-value ≤ 0.05. A minimum of n=3 data points per condition was required for each phosphopeptide and protein to be assessed for differential expression and statistical significance by limma and for downstream analyses. To retain maximum information without imputation or excluding IDs with any missing values, IDs with equal sample sizes per condition were grouped and subjected to differential expression analysis and multiple testing correction. ClusterProfiler^106^ v4.8.3 (with OrgDb from March 2023) and EnrichR^107^ (v3.2) were used for gene ontology term enrichment analysis, with p-values adjusted by the Benjamini–Hochberg procedure. Redundant terms were removed using the simplify function in ClusterProfiler. Custom backgrounds were made using the list of proteins in each mass spectrometry data from which the enriched IDs were extracted. For GO Biological Process, Molecular Function, and Cellular Component databases, v2023 was used. Oligodendrocyte-specific markers were retrieved from PanglaoDB^108^ and Tabula Muris^109^, and markers that had evidence in the literature to be expressed highly in other cell types of the brain from these datasets were filtered out. GSEA on protein expression data was done using the fgsea R package^110^, with the minimum size of the signature set at 20. For GO term enrichment analyses of significantly differentially expressed IDs (FC ≥ 1.5 and adj. p-val ≤ 0.05) and GSEA, all phosphoproteins and proteins that were detected at least three times across conditions being tested were used. Heatmaps represent only the IDs that had no missing values across all samples of given conditions presented.

The visualization of the kinome and phosphatome was done using Coral^111^ and CoralP^112^ in R. Only murine kinases and phosphatases with corresponding human homologs were included in the plot (kinases and phosphatases that only exist in mice and not human were not detected in the experiment). The visualization of STRING networks^113^ and the Enrichment analysis (ClueGo 2.5.9^114^) on signaling networks were done in Cytoscape (3.9.1)^115^ using Biological processes (2022.05.25) and WikiPathways (2022.02.23) databases. GO term with the lowest term p-value within the ontology group was shown. NicheNetR^27^ was used to evaluate ligand-receptor relationships between O4-negative (sender) and O4-positive cells (receiver) cells, aiming to identify those potentially regulating myelination in O4-positive cells cells. A signature associated with myelin maintenance was retrieved from MSigDB. O4-positive cells cell-specific proteins were defined as proteins that had log_2_(O4-positive cells/O4-negative cells) greater than 0 (adj. p-val ≥ 0.25), and O4-negative cell-specific proteins were defined as proteins that had log_2_(O4-positive cells/O4-negative cells) less than 0 in the protein expression data. All quantified phosphorylation sites by INSIGHT on receptors expressed by O4-positive cells were shown, irrespective of whether phosphorylation levels were higher in O4-positive or O4-negative cells. Thus, in cases where the receptors were more abundant in O4-positive cells but exhibited higher phosphorylation in O4-negative cells, suggestive of shared protein expression between the two populations, the phosphorylation levels were represented as negative values in the heatmap. The predictive power of each ligand in driving oligodendrocyte differentiation protein expression in the O4-positive cells relative to background protein expression was assessed to find ligands with the highest regulatory potential using NicheNetR. Receptors expressed in O4-positive cells that could bind the prioritized ligands from O4-negative cells and whose phosphorylation sites were mapped were then plotted.

To calculate the GBM module enrichment scores on protein expression data, the genes specific to each GBM module (neural-progenitor-like (NPC), oligodendrocyte-progenitor-like (OPC), astrocyte-like (AC), hypoxia-independent mesenchymal-like (MES1), hypoxia-dependent mesenchymal-like (MES2), G1/S, and G2/M cell cycle states) were retrieved from Neftel *et al*.^48^. Then, only the corresponding proteins that were detected and quantified by INSIGHT across all samples were used to calculate the enrichment scores using the single sample GSEA (ssGSEA) function from the GSVA package^116^. For the Sankey plot, the module that had the highest enrichment score in each tumor sample was plotted using networkD3.

To assess the synaptic proteins and phosphoproteins expressed by G6 and G12 PDX tumor cells, the SynGO annotations (version 20231201) were accessed and downloaded through the SynGO website^69^. Proteins (and phosphoproteins) with a fold change above 1.5 in primary tumors relative to normal cells were classified as expressed in primary tumor cells. Proteins (and phosphoproteins) without missing values in disseminated tumors were classified as expressed in disseminated tumor cells. For bulk RNAseq data, only the genes with log_2_(TPM+1) ≥ 0.1 were considered for SynGo analysis. To assess the overlap between the genes associated with metastasis, migration, invasion, epithelial-to-mesenchymal or mesenchymal-to-epithelial transition (EMT/MET), and synapse, lists of gene sets were curated from MSigDB using the msigdbr package (species = Homo Sapiens) and manually verified and curated using keywords (e.g., invasion), except for synapse-associated genes, which were downloaded from SynGO^69^.

To infer active kinases in the primary tumors, the pY, pS, and pT motifs (7 amino acid residues flanking the phosphorylated residues) were extracted from peptides that mapped to human proteome and analyzed through the Kinase Library^4,5^ with predetermined foreground (≥1.5-fold change, adj. p-val ≤0.1) and background sets (the rest of the phosphopeptide motifs that mapped to human proteome in the same experiment). For the inference of kinases that mediate the phosphorylation of specific residue sites, the Score Multiple Sites tab on the Kinase Library website was used. For all analyses, non-canonical kinases were excluded.

Various R packages for data processing, visualization, and plotting included paletteer (1.6.0), patchwork^117^ (1.3.0), magrittr (2.0.3), reshape2 (1.4.4), ggplotify (0.1.2), matrixStats (1.4.1), ggrepel (0.9.6), ggpubr (0.6.0), qdap (2.4.6), RColorBrewer (1.1-3), qdapTools (1.3.7), qdapRegex (0.7.8), qdapDictionaries (1.0.7), rlang (1.1.4), plyr (1.8.9), readxl (1.4.3), colorRamp2 (0.1.0), stringi (1.8.4), shadowtext (0.1.4), data.table (1.16.2), stringr (1.5.1), dplyr (1.1.4), purrr (1.0.2), readr (2.1.5), tidyr (1.3.1), tibble (3.2.1), tidyverse (2.0.0), ggplot2 (3.5.1), and ComplexHeatmap^118^ (2.18.0). complexupset^119^(1.3.3), ggVennDiagram^120^(1.3.0). Schematics were made using Biorender.com and Adobe Illustrator.

### Statistics and reproducibility

All statistical analyses were done using R (v4.3.1) in R studio. The statistical significance in plots of proteins and phosphopeptides identified by mass spectrometry are FDR-adjusted two-sided p-values (p’). Non-adjusted p-values were denoted as two-sided p only in the plots where quantifications were derived from LC-MS/MS. For co-immunofluorescence images, one-way ANOVA was used to test whether there were any overall differences among the cells in the core, margin, and left hemispheres, followed by Tukey’s Honest Significant Difference (HSD) test to perform two-sided pairwise comparisons between conditions and control the family-wise error rate. Plots show adjusted two-sided p-values between conditions, and asterisks in the top left corner represent the p-values from ANOVA (**** = p-value ≤ 0.0001, *** = p-value ≤ 0.001; ** = p-value ≤ 0.01; * = p-value ≤ 0.05, ns = p-value > 0.05). All box plots show the median, interquartile range (IQR, 25th to 75th percentile), and whiskers (up to 1.5 × IQR).

## Data and code availability

Raw MS files and TMT channel mapping information can be accessed on the PRIDE database (PXD056965). Codes used in this study are available at https://github.com/ryuhjinahn. The R image used for the RNA-seq data analysis is also available.

## Acknowledgment

We thank Connor Dobson, Beth Grace, Erin Rousseau, Claudia Varela, and Alex Jaeger for providing tissues for early pilot experiments. We thank Isabelle Carrier (Lady Davis Institute/McGill University) for the co-IF staining protocol, and Jason Godfrey, Bokai Song, and Weixi Kang for their critical reading of the manuscript and the White lab members for their feedback. We also thank the Division of Comparative Medicine (DCM), Kathleen Cormier from the Pathology Core facility, Glenn Paradis and Michele Griffin from the Flow cytometry Core facility, Stuart Levine, Gideon Donahue, and Avanyish Toniappa from the BioMicro Centre, and Jeffrey Kuhn from the Microscopy/Imaging Core for their tremendous guidance and training. We gratefully acknowledge John Martello for his generous philanthropic gift that supported part of this work. This work was also supported by NIH Grant CA283114, the Koch Institute Support Grant P30-CA14051 from the National Cancer Institute, and Cancer Center Support (core) Grant P30-CA14051 from the NCI to the Barbara K. Ostrom (1978) Bioinformatics and Computing Core Facility of the Swanson Biotechnology Center. RA was supported by the Next Generation of Scientists award (Grant 1143397) from the Cancer Research Society and the Ludwig Postdoctoral Fellowship from MIT Ludwig Center. ADD was supported by the National Science Foundation Graduate Research Fellowship. CTF was supported by a graduate fellowship from the Ludwig Center at MIT.

## Author contributions

RA and FMW conceived the study. RA, ADD, LL, YC, KKB, LLO, BLC, and JW performed all experiments. RA performed all mass spectrometry-associated sample processing, experiments, and data analyses. RA performed all data visualizations. DB, KKB, LLO, BLC, and JS generated all the PDX tumor materials. DB, AT, and JS provided support related to the PDX models. RA, TMY, and JLJ performed Kinase Library analyses. RA, INK, and CTF optimized the mass spectrometry instrument methods. GZ contributed to the development of computational workflow. CAW re-processed and analyzed published RNAseq raw data and wrote the associated method. JW performed confocal imaging. RA and FMW wrote the manuscript with feedback from all authors.

## SI Appendix

Extended Data Figures & Legends Supplementary Tables and Legends Supplementary Data Legends Supplementary Methods

## Extended Data Figure Legends

**Extended Data Figure 1.** Optimization of fixation method to preserve the phosphoproteome.

**Extended Data Figure 2.** Phosphoproteome and proteins remain intact after the INSIGHT workflow allowing for antibody or fluorescent endogenous protein-based cell isolation.

**Extended Data Figure 3.** Characterization of the G6 and G12 PDX tumor-bearing mouse brains.

**Extended Data Figure 4.** Phosphoproteome and protein expression within primary tumor cells and normal cells from the G12 PDX model.

**Extended Data Figure 5.** Expression of EGFR and pT693-EGFR in G6 PDX tumor cells.

**Extended Data Figure 6.** Expression of EGFR and pT693-EGFR in G12 PDX tumor cells.

**Extended Data Figure 7.** Signaling network analysis of G6 and G12 disseminated and primary tumor cells *in vivo*.

**Extended Data Figure 8.** Overlap between synapse and dissemination-associated gene signatures.

**Extended Data Figure 9.** Differentially expressed phosphopeptides and proteins in G6 and G12 primary and disseminated tumor cells *in vivo*.

**Extended Data Figure 10.** Proliferation-associated signaling is suppressed in disseminated tumor cells relative to primary tumor cells of G6 and G12 PDX models.

## Supplementary Table Legends

**Supplementary Table 1**. Differential expression between O4-positive and O4-negative cells

**Supplementary Table 2**. Differential expression between primary vs. normal and primary vs. disseminated in G6 and G12.

**Supplementary Table 3**. G6 and G12 PDX RNAseq counts.

**Supplementary Table 4**. Summary of clinically relevant characteristics of G6 and G12 PDX models

## Supplementary Data Legends

**Supplementary Figure 1.** Gating strategies for flow cytometry

